# AGS3-based optogenetic GDI induces GPCR-independent Gβγ signaling and macrophage migration

**DOI:** 10.1101/2024.06.04.597473

**Authors:** Waruna Thotamune, Sithurandi Ubeysinghe, Chathuri Rajarathna, Dinesh Kankanamge, Koshala Olupothage, Aditya Chandu, Bryan A. Copits, Ajith Karunarathne

## Abstract

G protein-coupled receptors (GPCRs) are efficient Guanine nucleotide exchange factors (GEFs) and exchange GDP to GTP on the Gα subunit of G protein heterotrimers in response to various extracellular stimuli, including neurotransmitters and light. GPCRs primarily broadcast signals through activated G proteins, GαGTP, and free Gβγ and are major disease drivers. Evidence shows that the ambient low threshold signaling required for cells is likely supplemented by signaling regulators such as non-GPCR GEFs and Guanine nucleotide Dissociation Inhibitors (GDIs). Activators of G protein Signaling 3 (AGS3) are recognized as a GDI involved in multiple health and disease-related processes. Nevertheless, understanding of AGS3 is limited, and no significant information is available on its structure-function relationship or signaling regulation in living cells. Here, we employed *in silico* structure-guided engineering of a novel optogenetic GDI, based on the AGS3’s G protein regulatory (GPR) motif, to understand its GDI activity and induce standalone Gβγ signaling in living cells on optical command. Our results demonstrate that plasma membrane recruitment of OptoGDI efficiently releases Gβγ, and its subcellular targeting generated localized PIP3 and triggered macrophage migration. Therefore, we propose OptoGDI as a powerful tool for optically dissecting GDI-mediated signaling pathways and triggering GPCR-independent Gβγ signaling in cells and *in vivo*.

## 1. Introduction

G protein-coupled receptors (GPCRs) represent ∼5% of the human genome, have immense pathophysiological significance, and thus a primary target for drug discovery (1–4). GPCRs constitute one of the largest and most diverse superfamilies of cell surface receptors that sense a wide array of extracellular ligands, ranging from photons to small molecules and peptides (5). Upon ligand binding, GPCRs activate the G-protein heterotrimer (Gαβγ) through GDP to GTP exchange on the Gα, which leads to GαGTP and Gβγ dissociation (6, 7). These G proteins then modulate downstream signaling, including activating kinases, lipases, GTPases, and ion channels, to support cellular life (8). The physiological outcomes of activated G proteins depend on the type of G protein coupling of the receptors and the Gβγ composition of the cell. For instance, heterotrimers with Gαq, 12/13 signals through PLC and RhoA pathways (9, 10), while Gαs and Gαi control the secondary messenger cAMP (11, 12). Although, all heterotrimeric pathways produce Gβγ, the signaling intensity of Gβγ is more prominent upon activation of Gαi/o and Gαs heterotrimer (13). Most cells show highly downregulated expression of Gαq, and thus, the concentration of Gβγ generated upon activation of this pathway is relatively minor (14). Though initially considered a passive signaling regulator of Gα activity, during the last two decades, Gβγ has been identified as a major signaling actuator that controls numerous effector pathways, including G protein-gated inwardly rectifying K^+^ channels, adenylyl cyclases, phospholipase C (PLC), phosphoinositide 3-kinase γ (PI3Kγ) and GPCR kinases (15–19).

Although GPCR-mediated activation of G protein heterotrimers is considered the major active regulator of numerous cellular activities in response to external stimulation (1, 20–23), it is understood that activities of the same downstream signaling components, albeit at lower intensities, are required for the ambient signaling of cells (24). Recent studies have indicated intricate molecular processes beyond the canonical GPCR-based model consisting of alternative forms (accessory proteins) of heterotrimeric G protein activity modulation independent of receptor activation (25, 26). Activators of G protein Signaling (AGS) family proteins are one such major regulators. Evidence indicates that AGS family members impose several distinct modes of G protein heterotrimer regulation and invoke GPCR-independent G protein signaling (27). AGS family proteins consist of three different classes of proteins based on their mechanism of action: (i) Class I AGS proteins mainly function as Guanine nucleotide exchange factors (GEFs) to induce GDP to GTP exchange and heterotrimer activation (like GPCRs), (ii) Class II AGS proteins function as Guanine nucleotide dissociation inhibitors (GDIs) as their specific G protein regulatory motifs (GPR) facilitate the binding to inactive GαiGDP form and repel Gβγ independent of GPCR activation, (iii) whereas Class III AGS proteins tend to interact with heterotrimers and promote effector activation, which is independent of the nucleotide exchange (28–30). As many G protein functions are evaluated by adding extracellular ligands for their respective GPCRs, the inability to use such a method to trigger and detect the action of AGS family proteins has been an obstacle to a clear understanding of their signaling and regulation. To overcome this, we used protein engineering to generate a novel OptoGDI capitalizing on the light-sensitive Cryptochrome 2 photolyase homology (CRY2PHR, aa 1-498) domain in combination with the more efficient consensus GPR motif derived from AGS3 (29, 31). Using OptoGDI, we show for the first time the real-time activity of a GDI protein in living cells. We also show the feasibility of using this approach to optically invoke standalone subcellular Gβγ signaling, including PI3K activation and directional cell migration. The presented data indicate that this novel optogenetic approach will allow spatiotemporal control over various crucial Gβγ mediated signaling processes, such as the regulation of ion channels and kinases independently of canonical GPCR signaling pathways (32–35), and therefore elucidate molecular underpinnings of complex regulatory networks governing cell signaling and behaviors. Such findings can potentially advance our understanding of fundamental cellular processes and offer new avenues for therapeutic intervention.

## 2. Results

### 2.1 Engineering of CRY2-based OptoGDIs

Our goals were to examine the regulatory role of AGS3 in signal processing in living cells and to create an optogenetic strategy to control standalone Gβγ signaling. The approach aimed to present an AGS3-derived Gα binding domain on optical command to the heterotrimer at the plasma membrane, releasing Gβγ. AGS3 is identified as a guanine nucleotide dissociation inhibitor (GDI) and possesses four G protein regulatory motifs (GPRs), which mediate its binding to the inactive Gαi/oGDP in the heterotrimer (29). The need for a significant excess Gβγ to disrupt the Gα-AGS3 complex indicated that AGS3 could release Gβγ without GDP to GTP exchange at Gα (36).

Four GPR motifs in AGS3 may enhance AGS3-heterotrimer reaction cross section and thus provide effective separation of Gβγ from GαGDP. A consensus GPR motif (GPRcn) was proposed by aligning the four GPR repeats from five species (37). GPRcn possesses an N-terminal negative charge from a *Glu* duo, followed by a hydrophobic cluster composed of *Phe* duo, *Leu* duo, and several C-terminal hydrophilic residues (*Asp-Asp-Gln-Arg*), and has been ten-fold effective in promoting heterotrimer dissociation than the Gβγ hotspot binding peptide, SIGK (38).

Consistent with the current predictions based on the primary sequence analysis, using *in silico* modeling, we show that the consensus sequence derived from the IVth GPR motif of AGS3 is likely to adopt an alpha-helical conformation (37, 39). Furthermore, our protein-protein docking data show a tight binding affinity between the modeled consensus GPR motif and GαiGDP, indicated by the several key interactions. Out of 30 potential configurations, more than 40% of configurations showed that the GPR helix binds to the same Gαi region. In particular, we found interactions between Thr1, Met2, Glu4, Phe7, Phe8, Leu10, Leu11, Ser14, Gln15, Arg18, and Met19 of the GPR with Arg205, Arg208, Ile212, Phe215, Glu216, Glu245, Lys248, Leu249, Asp251, Ser252, Asn256, Lys257, Trp258, Phe259, and Arg311 of GαiGDP, respectively. Based on our modeled structure, the GPR motif is likely to be an amphipathic helix (Fig. 1A). Using AGS3 full sequence and site-directed mutagenesis, we generated GPRcn, and fused it to the C-terminus of *Arabidopsis* CRY2-PHR tethered mCherry (31). Similar to the full wild-type AGS3 protein, CRY2-mCherry-GPRcn also showed a cytosolic distribution. Upon blue light exposure, CRY2PHR binds to CIBN(31). Since the goal was to generate Gβγ at the plasma membrane on an optical command, we generated N-terminally myristoylated/palmitoylated CIBN (Lyn-CIBN) using CIBN-CAAX (Fig.1B). Upon blue light illumination, CRY2-mCherry-GPRcn was recruited to the plasma membrane, indicating its heterodimerization with CIBN (Fig. 1B-right). To determine whether GPRcn interacts with plasma membrane-bound Gαi/oGDP and repel Gβγ, we employed the Gβγ9 translocation assay (13, 40). We have previously established GPCR activation-induced Gγ9 translocation (as Gβγ complex) from the plasma membrane to endomembranes as an assay to measure GPCR-G protein activity in living cells (40). As a positive control, we show the robust Gγ9 translocation exhibited by HeLa cells expressing Venus-Gγ9 upon activation of α2AR-CFP with 100 μM norepinephrine (Fig. 2A).

**Fig. 1:**
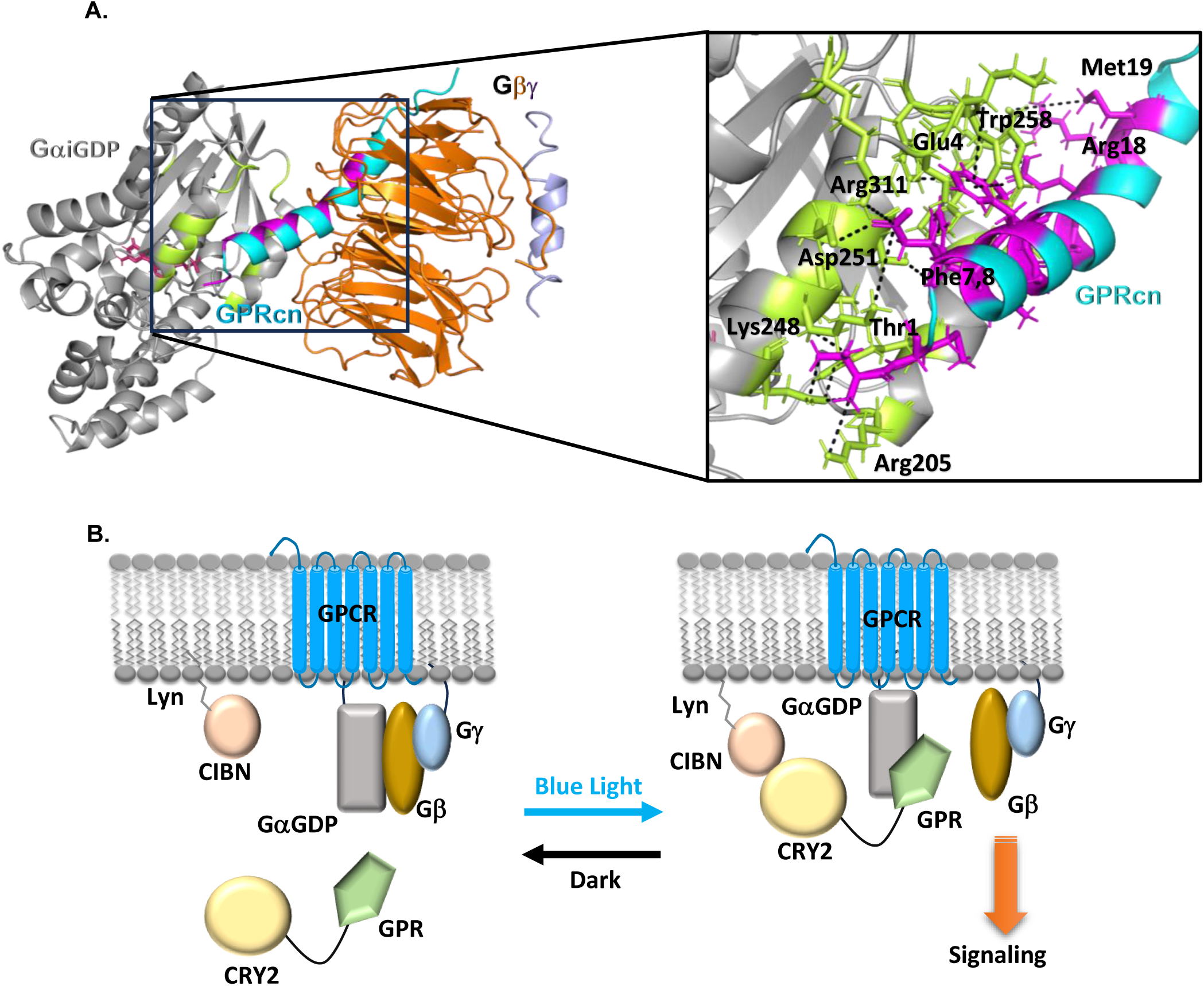
AGS3-GPR consensus peptide interacts with Gai subunit. **(A)** Diagram depicting the interactions between GαiGDP- Gβγ and GαiGDP-GPR consensus peptide. GαiGDP- gray, Gβ – orange, Gγ – light blue, GPR motif-cyan. GαiGDP residues that interact with GPR peptide are shown in green. GPR residues that interacts with GαiGDP are shown in pink. (PDB ID 7E9H and 2V4Z) **(B)** Scheme for optical recruitment of Cryptochrome 2 based GPR peptide to the plasma membrane.

**Fig. 2:**
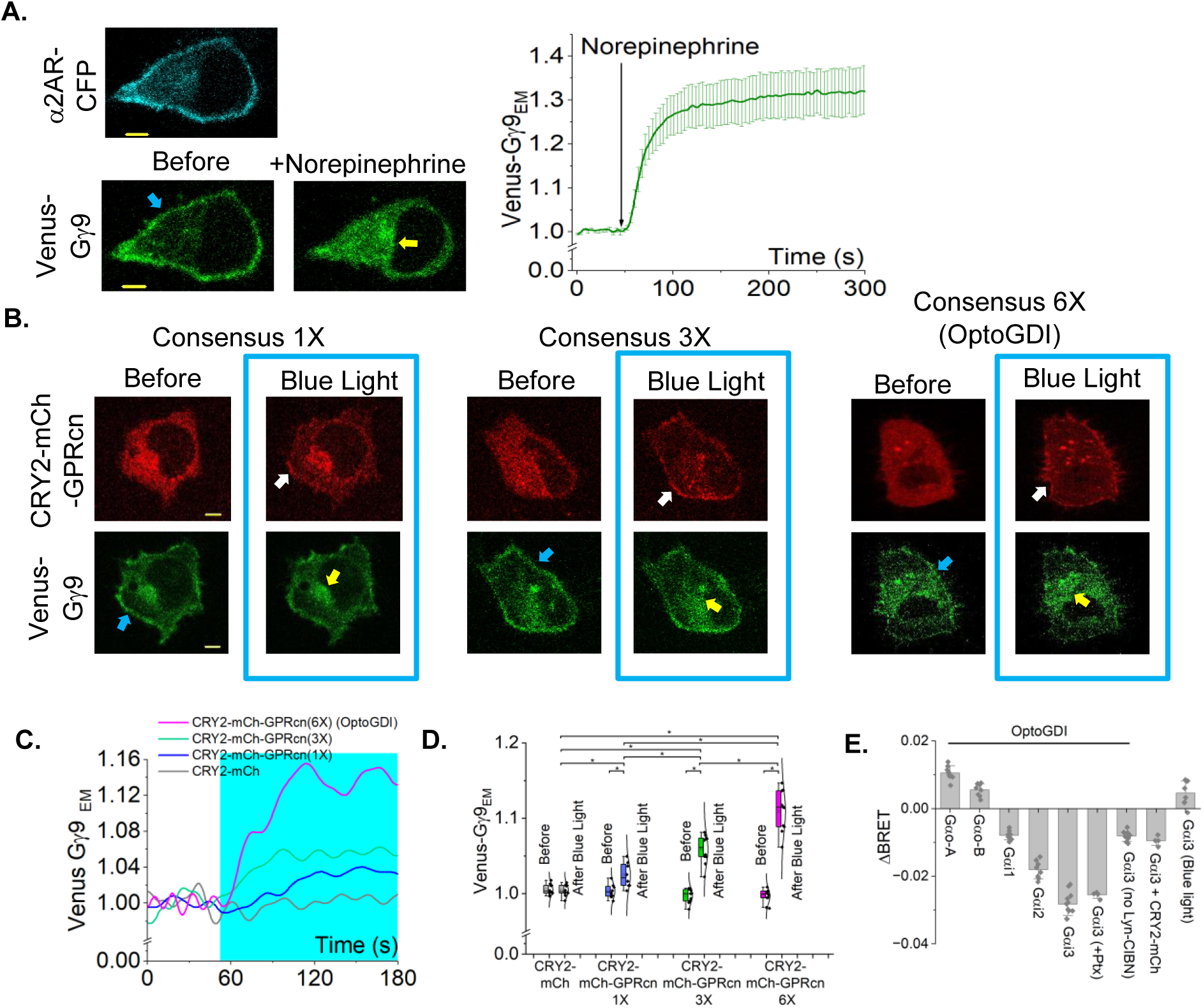
OptoGDI induces Gβγ translocation upon blue light exposure. **(A)** HeLa cells expressing α2AR- CFP and Venus-Gγ9 show robust Gβγ translocation upon 100 mM norepinephrine addition. The plot shows the baseline normalized Venus fluorescence in endomembranes over time. The accumulation of Gγ9 in internal membranes is indicated by yellow arrows. The Venus-Gγ9 loss from the plasma membrane is indicated by the blue arrow (n=10). **(B)** HeLa cells expressing Venus-Gγ9, Lyn-CIBN, and CRY2- mCh- GPRcn (1X), CRY2- mCh- GPRcn (3X), or CRY2- mCh- GPRcn (6X) (OptoGDI) show different extents of Venus-Gγ9 translocation to the endomembranes, upon blue light exposure. Yellow arrows indicate the accumulation of Gγ9 in internal membranes. The white arrow indicates the plasma membrane recruitment of CRY2-mCherry-GPRcn variants. The Venus-Gγ9 loss from the plasma membrane is indicated by the blue arrow. **(C)** The plot shows the baseline normalized Venus fluorescence in endomembranes over time. (n= 15 for 1X, n= 14 for 3X, n=17 for Opto-GDI). **(D)** The whisker box plot compares Gγ9 translocation extents to the basal Gγ9 fluorescence at endomembranes. Blue box indicates the blue light exposure. Average curves were plotted using cells from ≥3 independent experiments. **(E)** The bar chart showing ΔBRET between Rluc8 tagged Gα and GFP2 tagged γ9 under different experimental conditions. OptoGDI with Gα0-A (n=8), OptoGDI with Gα0-B (n=8) ), OptoGDI with Gαi1 (n=8), OptoGDI with Gαi2 (n=8), OptoGDI with Gαi3 (n=8), OptoGDI with Gαi3 and Ptx (n=4), OptoGDI with Gαi3 but no Lyn-CIBN (n=8), OptoGDI with CRY2-mCh (n=4), and Gαi3 with blue light (n=8). The error bars represent SD (standard deviation). The scale bar = 5 µm. CFP: cyan fluorescent protein; mCh: mCherry; EM: endomembranes; Ptx: Pertussis toxin.

We first examined the blue light-induced activity of CRY2-mCherry-GPRcn(1X), which only contains one GPRcn, in HeLa cells also expressing Lyn-CIBN and Venus-Gγ9. Upon blue light-induced plasma membrane recruitment of CRY2-mCherry-GPRcn(1X) (Fig. 2B-white arrow), cells showed a minor yet significant Gγ9 translocation (Fig. 2B-left panel, plot-blue curve, one-way ANOVA*: F*_1, 18_ = 60.49, *p* =<0.0001; Supplemental Table S1, A, and B) unlike GPCR activation-induced translocation (Fig. 2A). We believe an inefficient binding affinity for GαGDP and lack of proper spatial orientation for the interaction to be few of the several possible reasons underlying the optogenetic GDI-induced minor Gγ9 translocation. Since AGS3 consists of four GPR motifs at its C terminus, we hypothesized that increasing the number of consensus repeats in the optogenetic module may trigger a higher extent of Gβγ release from the heterotrimer. Therefore, we generated two new constructs, CRY2-mCherry-GPRcn(3X) and CRY2- mCherry-GPRcn(6X), in which we inserted flexible linker sequences between the consensus peptides.

Upon blue light exposure, CRY2-mCherry-GPRcn(3X) liberated a detectable amount of free Gβγ, while CRY2-mCherry-GPRcn(6X) exhibited significantly higher free Gβγ generation, measured using the change in endomembrane fluorescence due to Venus-Gγ9 translocation (Fig. 2B middle and right images, plot-green and pink curves). Additionally, we performed a similar experiment using CRY2PHR-mCherry, which does not contain a GPRcn motif at its C terminus to confirm that the observed Venus-Gγ9 is due to the Gβγ liberation from the GPRcn recruitment. Upon blue light-induced plasma membrane recruitment of CRY2-mCherry (Fig. S1A-top panel), cells did not show a detectable Gγ9 translocation (Fig. S1A-bottom panel, Fig. 2C and 2D-grey line and box). One-way ANOVA showed that CRY2-mCherry-GPRcn(6x) induces significantly higher Gβγ liberation than that of the other two versions (Fig. 2D-whisker box plot; one-way ANOVA*: F*_2, 24_ =25.0359, *p* =<0.0001; Supplemental Table S4, A, and B). Therefore, the studies henceforward are performed using CRY2-mCherry-GPRcn(6X), which we named OptoGDI. To investigate the Gαi/o subtype specificity of OptoGDI towards different heterotrimers, we used TRUPATH biosensors in which the heterotrimer dissociation can be detected using bioluminescence energy transfer between Rluc8 incorporated specific Gα type and GFP2-tagged Gγ9 (41). OptoGDI effectively generated free Gβγ from Gαi2 and Gαi3 heterotrimers upon exposure to blue light (Fig. 2E). However, OptoGDI could not liberate Gβγ from Gαi1, Gαo-A, and Gαo-B heterotrimers. The effect of OptoGDI on Gαi3 heterotrimer dissociation was not disturbed in cells treated overnight with pertussis toxin (Ptx) (200ng/mL), indicating such Gβγ generation is nucleotide exchange and Gαi-coupled GPCR independent. Control experiments with only OptoGDI showed minimum activity without its binding partner, Lyn-CIBN. Similarly, another control experiment with CRY2-mCh with Lyn-CIBN confirmed that the GDI portion is required for Gβγ liberation from Gαi-3 heterotrimers. Additionally, we did not observe any BRET signal after blue light stimulation of cells only expressing Gαi-3 heterotrimers without OptoGDI, confirming OptoGDI is required to liberate Gβγ.

### 2.2 OptoGDI-induced subcellular free Gβγ generation

Previous studies have shown the utility of several optogenetic tools to spatiotemporally control G protein signaling in subcellular locations upon light stimulations.(42–45) Therefore, next, to examine OptoGDI-induced subcellular G-protein signaling, we examined the ability of OptoGDI to induce subcellular Gβγ release and subsequent signaling. In HeLa cells expressing Lyn-CIBN, OptoGDI, and Venus-Gγ9, OptoGDI showed cytosolic, and Venus-Gγ9 displayed primarily plasma membrane distribution (Fig. 3A-left panel). We then exposed a selected sub-plasma membrane region to blue light using a localized blue light stimulus (445 nm, 26.3 μW/mm^2^) (Fig. 3A-middle panel-blue box). Robust recruitment of OptoGDI to the blue light exposed region of the cell and subsequent localized loss of Venus-Gγ9 membrane fluorescence, indicating Gβγ translocation (Fig. 3A-middle panel). Upon termination of the blue light, OptoGDI dissociation from the plasma membrane and Gγ9 recovery was observed, suggesting that OptoGDI induced a reversible heterotrimer dissociation (Fig. 3 A-right panel, B-Kymographs, and the plot). Additionally, the kymograph shows OptoGDI and Venus-Gγ9 localizations at the plasma membrane are inversely synchronized (Fig. 3B-kymographs and plot).

**Fig. 3:**
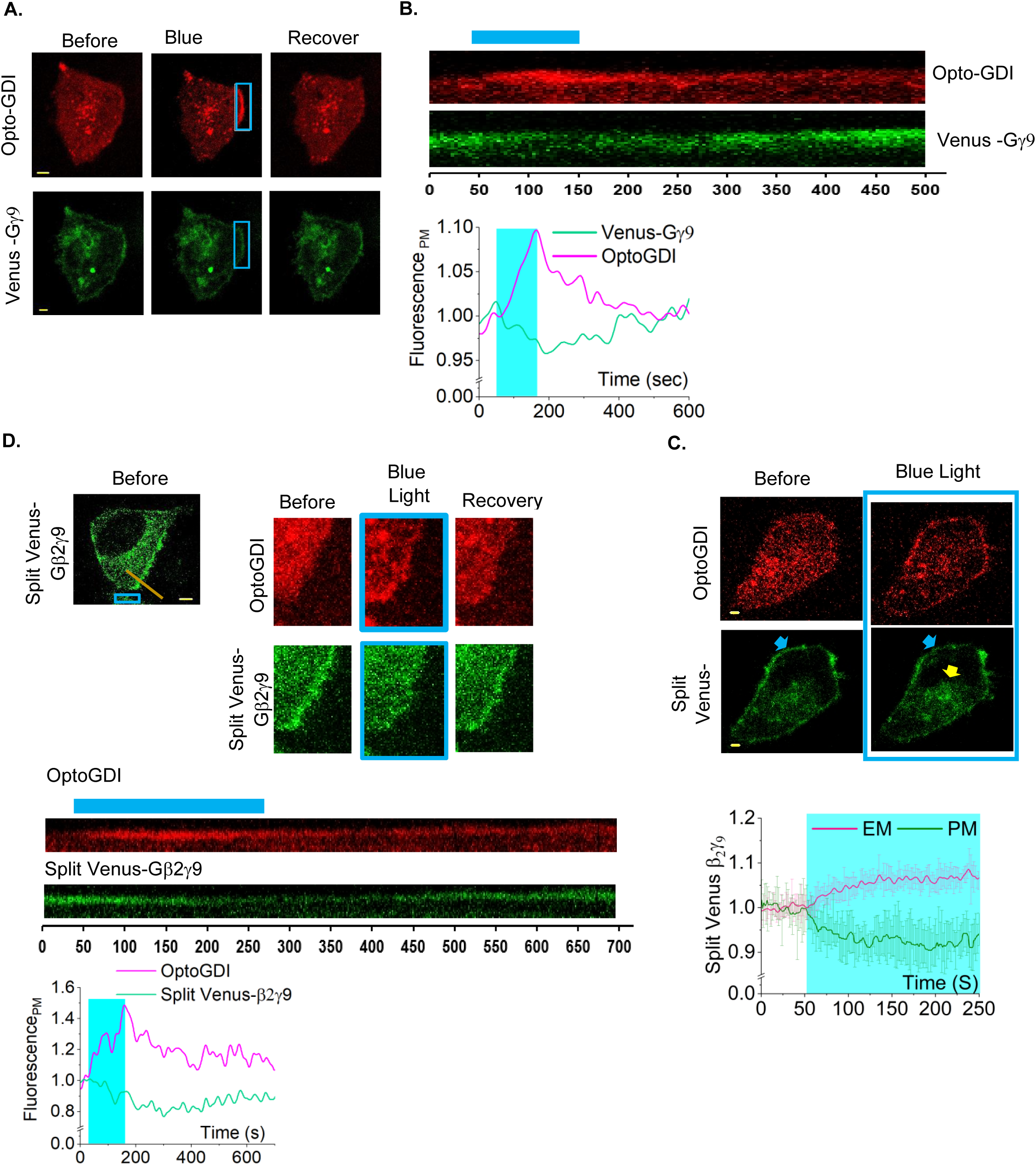
OptoGDI induces subcellular free Gβγ generation upon blue light exposure. (**A**) confined membrane region of a HeLa cell expressing OptoGDI, Lyn-CIBN, and Venus-Gγ9 was exposed to blue light. The cell exhibited a OptoGDI recruitment and Gγ9 translocation only on the blue light exposed region of the cell (n=12). **(B**) The kymographs and the plot showing the blue light induced OptoGDI recruitment and the Gγ9 loss and the recovery on the plasma membrane. **(C**) HeLa cells expressing Opto-GDI, Lyn-CIBN, and Split Venus-Gβ2γ9 show detectable free Gβγ translocation upon blue light exposure. The white arrow at the membrane indicates the OptoGDI recruitment after blue light stimulation. The blue arrow indicates Split Venus-Gβ2γ9 fluorescence loss from the plasma membrane. The plot shows the baseline normalized Venus fluorescence in endomembranes and the plasma membrane over time (n=8). **(D**) HeLa cells expressing OptoGDI, Lyn-CIBN, and Split Venus-Gβ2γ9 show subcellular free Gβγ generation upon blue light exposure to a confined membrane region. Kymographic view of the respective region of the cell shows OptoGDI and Split Venus-Gβ2γ9 dynamics at the plasma membrane over time. The orange line on the cell image indicates the membrane region used to create the kymograph. (n=12). Yellow arrows indicate the Venus Fluorescence increase on the plasma membrane when Opto-GDI is cytosolic. Average curves were plotted using cells from ≥3 independent experiments. The blue box indicates blue light exposure. The error bars represent SD (standard deviation of mean). The scale bar = 5 µm.

To ascertain that the observed Gγ9 translocation upon OptoGDI recruitment to the plasma membrane is not an experimental artifact and truly due to Gβγ liberation from the heterotrimer, we used a fluorescent complementation approach to track the distribution of both Gβ and Gγ. We expressed OptoGDI, Lyn-CIBN, Venus (1–155)-γ9, and Venus (155–209)-β2 in HeLa cells (46). It has been shown that the two proteins must be proximal enough to assemble the fluorescence protein Venus (47). Venus (1–155)-γ9 and Venus (156–209)-β2 do not exhibit any fluorescence when they are not interacting(48). HeLa cells expressing above constructs combination showed a Venus fluorescence with a reasonable plasma membrane localization (Fig. 3C-bottom images and 3D), while OptoGDI was observed in the cytosol. Next, we recruited OptoGDI to the plasma membrane by exposing cells to blue light (Fig. 3C-top images-blue arrow).

Venus fluorescence on the plasma membrane was reduced (Fig. 3C-white arrow) with a corresponding fluorescence increase in endomembranes (Fig. 3C-yellow arrow and the plot). In the Gβγ dimer, the N-termini of Gβ and Gγ are only a few angstroms apart; therefore, the venus fluorescence appearance and OptoGDI induced translocation shows the complementation between Venus fluorescence protein fragments in Gβ and Gγ. Next, we also examined the feasibility of liberating subcellular free Gβγ by using split-venus Gβγ described above. Similar to Fig. 3A, we exposed a confined region of the plasma membrane to blue light and recruited OptoGDI (Fig. 3D). Unsurprisingly, we observed the loss of Venus fluorescence at the blue light-exposed plasma membrane region, confirming Gβγ liberation from the heterotrimer. Upon blue light termination, OptoGDI gradually returned to the cytosol, and a synchronized reverse Gβγ translocation back to the plasma membrane was observed, indicated by the venus fluorescence recovery. The images and kymograph clearly indicate that the Gβγ release and recovery are governed by OptoGDI concentration at the plasma membrane (Fig. 3D-cell images and kymograph). This data also indicated that OptoGDI-induced Gβγ release is reversible and can be terminated by stopping blue light.

### 2.3 OptoGDI-liberated Gβγ modulates PLC**β**-induced PIP2 hydrolysis

Gq-coupled GPCR activation induces a self-attenuating PIP2 hydrolysis (10). We showed that the observed PIP2 hydrolysis attenuation is due to the loss of Gβγ from the highly effective lipase complex, GαqGTP-PLCβ-Gβγ through the activation of PLCβ and generation of the less effective lipase GαqGTP-PLCβ (10). We also showed that PIP2 hydrolysis can be rescued by providing Gβγ to a system with PIP2 hydrolysis partially attenuated due to the loss of Gβγ from GαqGTP-PLCβ-Gβγ (10). Here, we first examined whether Gβγ generated by membrane recruited-OptoGDI retards PIP2 hydrolysis attenuation in HeLa cells expressing GRPR (a Gq-coupled GPCR), Venus-PH (PIP2 sensor), Lyn-CIBN, and OptoGDI upon activation of GRPR using 1 µM bombesin. After PIP2 hydrolysis reached equilibrium, we exposed cells to blue light and recruited OptoGDI to the plasma membrane. However, cells didn’t exhibit the expected PIP2 hydrolysis rescue that occurs due to free Gβγ generation (Fig. 4A). To confirm that free Gβγ can rescue the attenuated PIP2 hydrolysis under the same experimental conditions, we examined the PIP2 hydrolysis rescue in HeLa cells expressing GRPR, α2AR-CFP, and Venus-PH, and PIP2 hydrolysis was partially attenuated after GRPR activation (Fig. 4B). The cells exhibited the characteristic PIP2 hydrolysis upon activation of GRPR using 1 μM bombesin. After the PIP2 recovery reached the steady state, activation of α2AR using 100 μM norepinephrine-induced significant PIP2 re-hydrolysis (Fig. 4B), indicating the α2AR activation can induce PIP2 hydrolysis rescue.

**Fig. 4:**
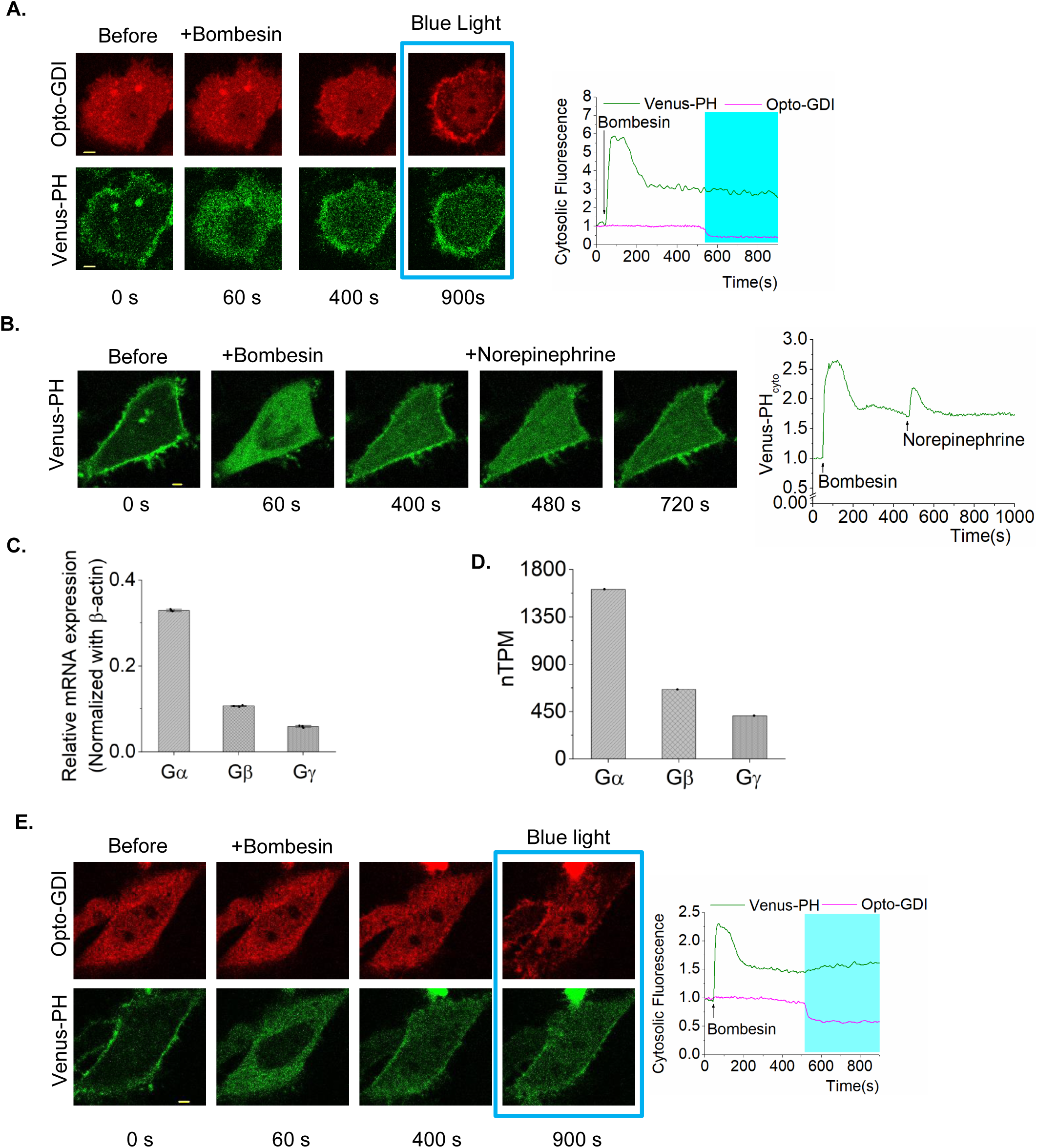
OptoGDI induces secondary PIP2 hydrolysis in Gq-GPCR activated cells. **(A)** HeLa cells expressing GRPR, Venus-PH, OptoGDI and Lyn-CIBN exhibited robust PIP2 hydrolysis upon 1 μM bombesin addition. OptoGDI was then recruited to the plasma membrane after the PIP2 hydrolysis reached equilibrium. The cells didn’t exhibit a rescue of PIP2 hydrolysis (n=15). **(B)** GRPR in HeLa cells expressing α2AR-CFP and Venus-PH was activated using 1 μM bombesin and the cells exhibited robust PIP2 hydrolysis. α2AR was then activated using 100 μM norepinephrine after the PIP2 hydrolysis reached the equilibrium. The cells exhibited robust secondary PIP2 hydrolysis indicating the PIP2 hydrolysis rescue (n=10). **(C)** Normalized relative mRNA expression level from RNAseq data of HeLa cells showing elevated expression of endogenous G-protein α compared to G-protein β and γ, normalized to the mRNA expression level of β-actin. (n=3) **(D)** Normalized transcripts per million (nTPM) levels of human G-protein α, β and γ levels from Human protein Atlas showing the elevated level of G-protein α, compared to the G-protein β and γ. **(E)** HeLa cells expressing GRPR, Venus-PH, Gβ1, Lyn-CIBN and OptoGDI were exposed to 1 μM bombesin. OptoGDI was then recruited to the plasma membrane of the cell after the PIP2 hydrolysis reached equilibrium. The cell showed detectable secondary PIP2 hydrolysis as indicated by the Venus-PH translocation to the cytosol. (n=12). The scale bar = 5 µm. Blue box indicates the blue light exposure. GRPR: Gastrin Releasing Peptide receptor; PIP2: Phosphatidylinositol 4,5-bisphosphate; PM: plasma membrane; PH: Pleckstrin Homology; Cyto: cytosolic fluorescence.

Compared to Gβγ dimers released upon Gi/o coupled GPCR activation (Fig. 2A), OptoGDI liberated only a fraction (Fig. 2B). To confirm that the limited free Gβγ generation is the underlying reason behind the lack of OptoGDI-induced PIP2 hydrolysis rescue (Fig. 4A), compared to the Gi/o coupled GPCR induced (Fig. 4B), we explored avenues to increase the concentration of liberated Gβγ by OptoGDI. For this, when we examined the endogenous G-protein expression levels in HeLa cells using HeLa cells RNAseq data, compared to the Gβγ dimers, Gα showed a higher expression (Fig. 4C). Data from human protein atlas also indicated a similar trend (Fig. 4D). Therefore, we hypothesized that a fraction of liberated Gβγ induced by OptoGDI is sequestered by the excess Gα subunits available on the plasma membrane. Our previous computation modeling also suggested a similar regulatory mechanism (13). Therefore, to reduce free GαGDP availability, we additionally expressed Gβ in HeLa cells, also expressing GRPR, Venus-PH, OptoGDI, and Lyn-CIBN. Upon GRPR activation, the cells showed robust PIP2 hydrolysis, which self-attenuated, reaching a steady state. Upon blue light induced OptoGDI recruitment to the plasma membrane, a detectable rescue of PIP2 hydrolysis was observed, indicating that free Gβγ at the plasma membrane can enhance the lipase activity of PLCβ (Fig. 4E). In a similar experiment, where we expressed CRY2-mCherry in the place of OptoGDI, blue light exposure failed to rescue PIP2 hydrolysis (Fig. S2A).

We next examined whether Gβγ generated by OptoGDI could reduce the rate of PIP2 hydrolysis attenuation that occurred after initial Gq-GPCR activation. Similar to the previous experiment, we examined GRPR-induced PIP2 hydrolysis in HeLa cells expressing GRPR, Venus-PH, Lyn-CIBN, and either CRY2-mCherry (control) or OptoGDI upon activation of GRPR using 1 µM bombesin. Here, we employed two controls. In the first control, we examined PIP2 dynamics in cells where we recruited CRY2-mCherry to the plasma membrane using blue light exposure after PIP2 hydrolysis reached the maximum (Fig. 5A-top panel images and 5B plot-green curve). We then examined PIP2 dynamics without OptoGDI recruitment to the plasma membrane in the second control. Interestingly, both the controls showed similar PIP2 dynamics (Fig. 5A-middle panel, 5B and 5C). Blue light-induced OptoGDI recruitment to the plasma membrane after the maximum PIP2 hydrolysis significantly retarded the PIP2 hydrolysis attenuation compared to the control cells (Fig. 5A-bottom panel and 5B-blue curve). The whisker box plot shows that the rate of PIP2 hydrolysis attenuation is significantly reduced in cells with membrane-recruited OptoGDI (Fig. 5C, one-way ANOVA*: F*_2, 28_ =66.825, *p* =2.188E-11; Supplemental Table S5, A, and B). Here, we believe that OptoGDI recruitment to the plasma membrane of GRPR-activated cells provides additional Gβγ to retard the PIP2 hydrolysis attenuation due to the reaction GαqGTP-PLCβ-Gβγ ⇌ GαqGTP-PLCβ + Gβγ, in which Gβγ is lost due to translocation (10). Therefore, we propose that OptoGDI pushes this reaction left, creating a more efficient lipase, and reducing the PIP2 hydrolysis attenuation (Fig. 5D).

**Fig. 5:**
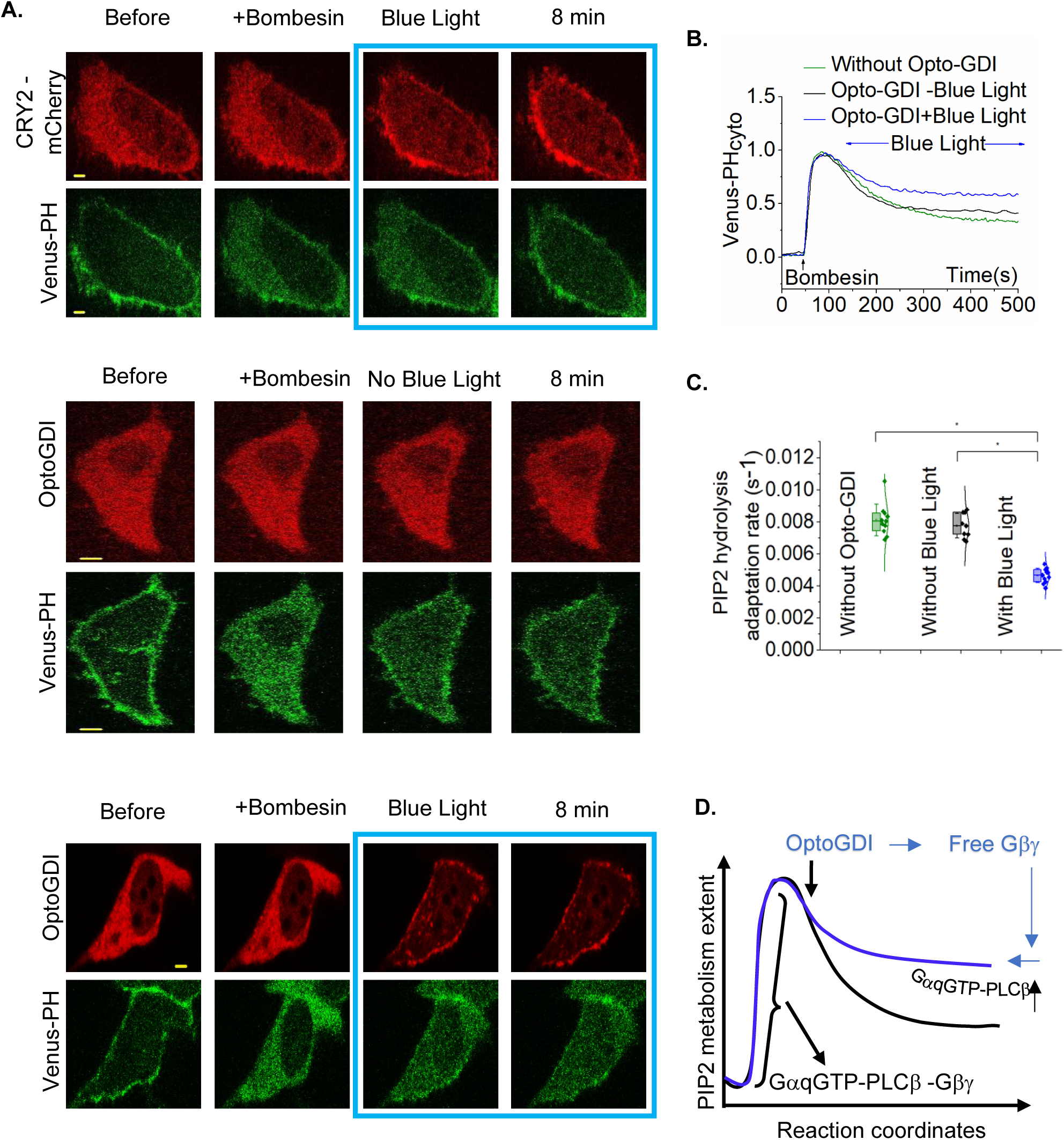
OptoGDI modulates PLCβ-induced PIP2 hydrolysis. **(A)** Top panel-GRPR, Venus-PH, Lyn-CIBN and CRY2-mCh were expressed in HeLa cells and cells were exposed to bombesin first and then exposed to blue light after the PIP2 hydrolysis reached the maximum. CRY2-mCh membrane recruited cells exhibited the typical PIP2 hydrolysis response followed by a PIP2 hydrolysis attenuation (n =11). Middle panel-HeLa cells expressing GRPR, Venus –PH, Lyn-CIBN, and OptoGDI showed robust PIP2 hydrolysis upon bombesin addition. The PIP2 hydrolysis attenuation rates were also similar to the previous condition (n=9). Bottom panel-HeLa cells expressing GRPR, Venus-PH, Lyn-CIBN and OptoGDI were first exposed to bombesin and then exposed to blue light and Opto-GDI was recruited to the plasma membrane when the PIP2 hydrolysis reached the maximum. The cells showed a robust PIP2 hydrolysis upon bombesin addition. The PIP2 hydrolysis attenuation rates were significantly lower than the control cells (n=11). **(B)** The corresponding plot shows the PIP2 sensor dynamics in the cytosol of the cells. **(C**) The whisker box plot shows the statistical differences in PIP2 hydrolysis attenuation rates between the control cells and OptoGDI recruited cells. **(D**) The schematic representation shows the mechanism of PIP2 hydrolysis attenuation. The scale bar = 5 µm. Blue box indicates the blue light exposure. GRPR: Gastrin Releasing Peptide receptor; mCh: mCherry fluorescent protein; PIP2: Phosphatidylinositol 4,5-bisphosphate; Cyto: cytosolic fluorescence; PH: Pleckstrin Homology.

### 2.4 Localized OptoGDI induces subcellular PIP3 generation and macrophage migration

Asymmetric activation of Gi/o GPCRs due to a chemokine gradient induces directional cell migration (49, 50). We have shown that Gβγ signaling at the leading edge, primarily Gβγ-induced PI3Kγ activation, generates traction forces through PIP3-mediated cytoskeleton remodeling and actively triggers the retraction of the trailing edge, facilitating the relocation of the cell body toward the chemokine gradient or the asymmetric optical stimulation (43, 51, 52). Here, we examined whether asymmetric GDI activity across a cell could trigger similar cell migration.

We expressed OptoGDI, Akt-PH-Venus (PIP3 sensor), and Lyn-CIBN, in RAW264.7 cells and then exposed a confined region of the cells to blue light (Fig. 6A top, blue box, Movie S1), and cells not only showed OptoGDI recruitment to the blue light exposed side of the cells but also an accompanying localized PIP3 generation (Fig. 6A-bottom, Movie S2). The kymographic views of the cells clearly show the blue light-induced plasma membrane-localized OptoGDI, PIP3 generation, and the subsequent movement of the cell towards blue light (Fig. 6B). We used CRY2-mCherry as a negative control to confirm that our observations were due to membrane recruitment of GPRcn motifs, however not because of CRY2 recruitment to the plasma membrane. We expressed CRY2-mCherry, Akt-PH-Venus, and Lyn-CIBN in RAW264.7 cells and examined the PIP3 generation and cell migration using blue light-induced localized CRY2-mCherry recruitment. As expected, localized CRY2-mCherry recruitment to the plasma membrane (Fig. S3A-top-blue box, Kymograph-top) did not show any detectable PIP3 generation or cell migration (Fig. S3A-bottom-blue box, Kymograph-bottom).

**Fig. 6:**
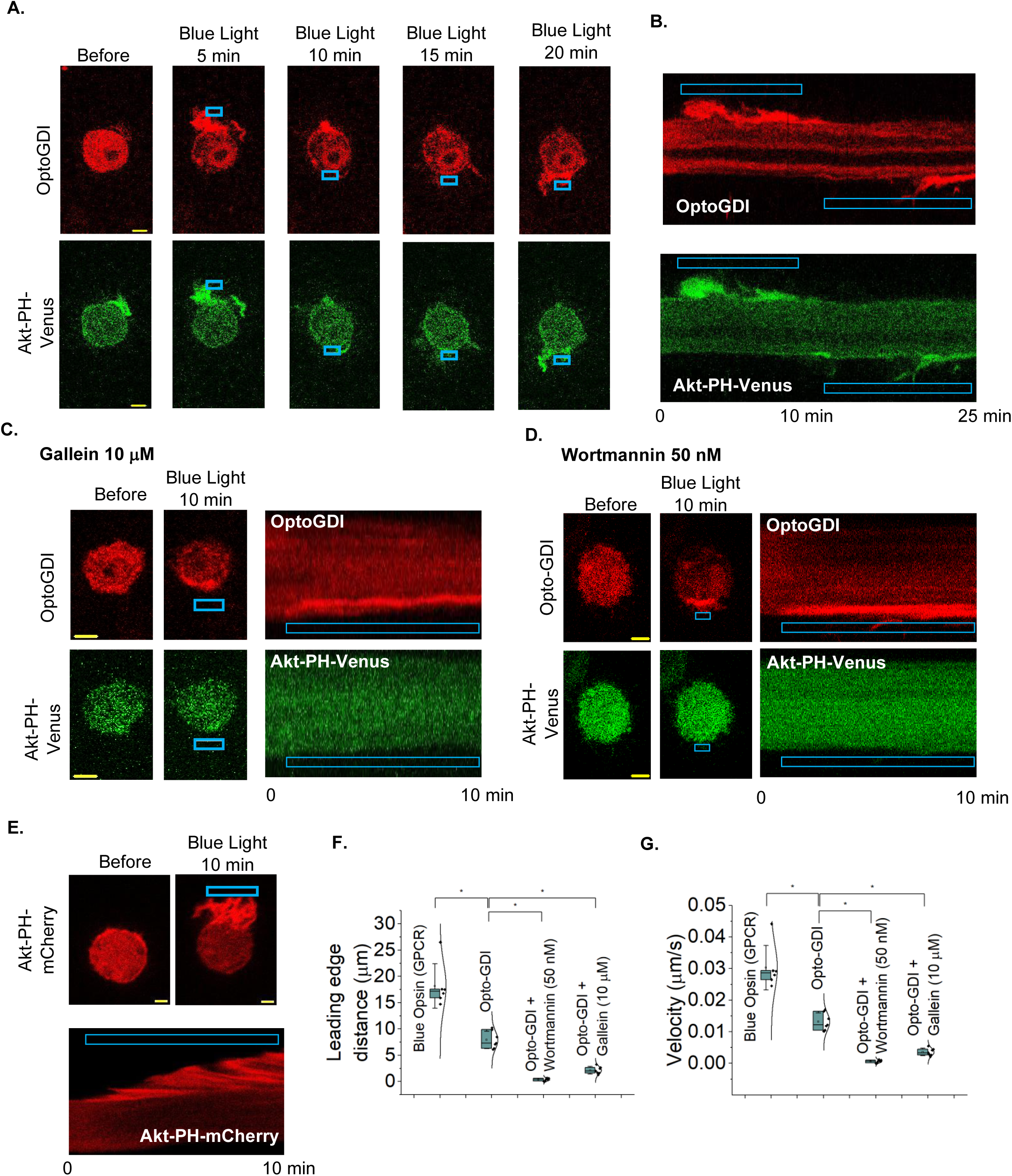
OptoGDI-triggered subcellular Gβγ induced PIP3 generation, and macrophage migration. **(A)** RAW cells expressing OptoGDI, Lyn-CIBN, and AKT-PH-Venus were optically activated to recruit OptoGDI to a confined region (top). The cell shows localized PIP3 production (bottom panel images) and the optogenetic activation of PIP3 production resulted in a detectable cell migration response towards the blue light. The blue box indicates the photoactivation (n=10). **(B)** Kymographs of the same cell show the accumulation of OptoGDI (red) and Akt-PH-Venus (green) in the leading edge of the migrating cell. **(C)** RAW 264.7 cells transfected with OptoGDI, Lyn-CIBN, and Akt-PH-Venus were subjected to localized optical activation in the presence of 10 μM gallein (Gβγ inhibitor). Cells were incubated with the inhibitor for 30 min at 37 °C before imaging and optical activation. The cells did not show a detectable migration or PIP3 generation. Kymographs of the same cell show the accumulation of OptoGDI (red) in the leading edge of a cell but no Akt-PH-Venus (green) accumulation (n=10). The blue box indicates photoactivation. **(D)** RAW 264.7 cells transfected with OptoGDI, Lyn-CIBN, and Akt-PH-Venus were subjected to localized optical activation in the presence of 50 nM wortmannin (PI3K inhibitor). Cells were incubated with the inhibitor for 30 min at 37 °C before imaging and optical activation. The cells did not show a detectable migration or PIP3 generation. Kymographs of the same cell show the accumulation of OptoGDI (red) in the leading edge of the cell but no detectable accumulation of the Akt-PH-Venus (green). The blue box indicates the photoactivation (n=12). **(E)** RAW 264.7 cells transfected with Blue Opsin and AKT-PH-mCherry were subjected to localized optical activation with confined blue light (Blue box), after the 50 mM 11-cis-retinal treatment. The cell showed an optogenetic activation induced localized PIP3 production and robust migration response towards the blue light. The kymograph of the same cell shows the accumulation of Akt-PH-mCherry (red) at the leading edge of the cell, indicating the PIP3 generation (n=9). **(F)**The whisker plot shows the extent of movement of the peripheries of the leading edge of control and pharmacologically perturbed RAW 264.7 cells in C-D. **(G)** The whisker plot shows the extent of migration velocities of the peripheries of the leading edge of the RAW cells in A-E. Average curves were plotted using cells from ≥3 independent experiments. The error bars represent SD (standard deviation of the mean). The scale bar = 5 µm.

As control experiments to ensure that the observed PIP3 generation is Gβγ mediated, and cell migration is PI3K activation-dependent, we performed a similar experiment, however either with cells exposed to 10 µM gallein; a Gβγ inhibitor (**43**), or 50 nM wortmannin, a PI3K inhibitor (**43**). In both control experiments, although the localized blue light induced subcellular OptoGDI recruitment, cells neither exhibited PIP3 generation nor directional cell migration (Fig. 6C and D). The OptoGDI-induced RAW264.7 cell migration extent and velocities observed in gallein, and wortmannin-treated cells were also significantly lower than in the untreated control cells (Fig. 5E and F, one-way ANOVA*: F*_1, 11_=120.349, *p* =<0.0001; Supplemental Table S10, A, and B and one-way ANOVA*: F*_1, 12_ =71.587, *p* =<0.0001; Supplemental Table S11, A, and B). As a positive control, we next compared the observed OptoGDI-induced RAW264.7 cell migration with blue opsin, a Gi/o-coupled GPCR, activation-induced macrophage migration. We expressed blue-opsin and Akt-PH-mCherry in RAW264.7 cells, which are exposed to 500 nM 11-cis-retinal before exposing cells to localized blue light activate opsins in one side of the cell. The cells showed robust migration towards blue light while forming lamellipodia at the activated leading edge of the cell (Fig. 6C cell images and kymograph). Upon blue opsin activation, Akt-PH-mCherry accumulation at the leading edge of the cell further indicated the robust PIP3 generation. One-way ANOVA indicated that the OptoGDI-induced RAW264.7 cells migration (distances and the migration velocities) is significantly lower than that of blue opsin-induced (Fig. 6E and F, one-way ANOVA*: F*_1, 11_ =35.346, *p* =<0.0001; Supplemental Table S6, A, and B and one-way ANOVA*: F*_1, 11_ =35.346, *p* =<0.00011; Supplemental Table S9, A, and B). This data collectively indicates that, though less potent than GPCR-induced, OptoGDI-triggered subcellular Gβγ can induce significant localized PIP3 generation and directional macrophage migration independent of GPCR activation.

## 3. Discussion

In this study, we demonstrate *in silico* structure-guided engineering of optogenetic GDI utilizing the AGS3 consensus GPR motif, enabling precise optical control of GDI-heterotrimer interactions to release signaling active Gβγ. Building on our previous work highlighting the substantial involvement of free Gβγ in PIP2 hydrolysis and its attenuation (10), we here demonstrated a negative impact on the canonical PIP2 hydrolysis attenuation observed upon Gq-GPCR activation through the OptoGDI-induced optogenetic release of free Gβγ. Using a Gγ subunit that provides the Gβγ complex the lowest observed membrane affinity, Gγ9 (53, 54), we show the reversible Gβγ release in single cells and from subcellular regions. To our understanding, this is the first direct demonstration of real-time GDI signaling in living cells. Furthermore, we show that spatially and temporally precise optical recruitment of OptoGDI to subcellular regions of the plasma membrane effectively initiates Gβγ-mediated signaling events, including localized PIP3 generation and associated macrophage migration. Although there are many studies on Gi/o-GPCR activation-induced and primarily Gβγ-mediated cell migration, the work presented here shows asymmetric standalone Gβγ signaling is sufficient to induce effective directional cell migration. As expected from a constitutively active signaling regulator, though OptoGDI-induced Gβγ generation and subsequent signaling were relatively lower, but they were significant. Although the observed Gβγ recovery at the plasma membrane upon ceasing blue light exposure was difficult to detect in single cells, which is a clear observation in GPCR activation-mediated signaling; the signaling reversibility is clear in subcellular OptoGDI signaling activation and termination. The reversibility of the PIP3 generation and migration direction observed upon reversing subcellular plasma membrane recruitment of OptoGDI indicated the ability to switch ON-OFF OptoGDI signaling on optical command.

The influence of AGS3 on a variety of signaling processes has been demonstrated. Among them, AGS3-influenced protein trafficking to the plasma membranes (55), μ-Opioid receptor-induced PKA signaling in primary striatal neurons (56), and the phosphorylation of cyclic AMP response element-binding protein (p-CREB) associated anti-apoptotic effects (57) are significant. The ability of the GPR motif of AGS3 to bind the G-protein heterotrimers, specifically, GαGDP through the upstream negatively charged and hydrophobic residues of the GPR motif liberating Gβγ has been suggested as the primary underlying molecular reasoning for AGS3 signaling (37). Also, it has been observed that the Gαi coupled GPCRs do not efficiently couple to GPR-bound GαiGDP, suggesting that, although the receptor interacts with the GαiGDP-GPR complex, this interaction stabilizes a receptor conformation that has a low affinity for agonist (37). It is also unclear whether GPR motif binding to GαGDP of the heterotrimer always results in free Gβγ release, or whether GPCRs can activate GPR-bound heterotrimers. Further, whether AGS3-heterotrimer interaction is a negative regulator for GPCR-G protein signaling and whether the GαiGDP-GPR complex acts as an active signaling entity are also unclear. Not only does the present study shed some light on these questions, but it also has provided a valuable avenue to understand AGS3 especially, GDI signaling in general, as well as a subcellular-amenable optogenetic tool to spatially and temporally control standalone Gβγ signaling.

Collectively, our *in silico* structural analysis indicates the GPR motif binding the GαGDP-Gβγ interface, while the provided single cell and subcellular experimental data show that OptoGDI induces the release of signaling active Gβγ, providing a molecular picture and mechanistic basis underlying AGS3 signaling. We showed the unique capabilities of a novel optogenetic GDI for spatially and temporally precise release of Gβγ in specific subcellular plasma membrane regions of living cells, thereby initiating a localized key signaling event in PIP3 generation, and a crucial physiological response leading to cell migration. Our finding not only provides a molecular description underlying AGS3 signaling but also demonstrates the feasibility of using GDIs to optically control Gβγ signaling independent of GPCR and Gα. The endowed precise spatial and temporal control can have a profound impact on the interrogation of Gβγ signaling in living cells and animals, exposing aberrant signaling implicated in various diseases, such as cancer and neurological diseases (58–60). By harnessing the power of optogenetics, we are not only expanding our understanding of the signaling of a molecule that we only know very little about but also providing a methodological template for *in silico*-guided engineering of optogenetic tools for signaling control.

## 4. Methods and Materials

### 4.1 Reagents

Reagent sources: Norepinephrine (Sigma Aldrich, St. Louis, MO), Gallein (TCI AMERICA), Wortmannin and Bombesin (Cayman Chemical, Ann Arbor, MI), 11-*cis*-retinal (National Eye Institute, Bethesda, MD). All reagents except 11-*cis*-retinal (ethanol) were initially dissolved in DMSO and then diluted in HBSS or cell culture medium before adding to cells.

### 4.2 DNA constructs

DNA constructs: Blue opsin (42) has been described previously (61). α2AR-CFP, Venus-Gγ9, Split Venus β2γ9, Akt-PH-Venus, Akt-PH-mCh, Venus-PH, CRY2-mCh, and CRY2-mCh-GPRcn constructs were kindly provided by the laboratory of Prof. N. Gautam at Washington University School of Medicine, St. Louis, MO. GRPR was a kind gift from the laboratory of Zhou-Feng Chen (62). The parent construct of AGS3 was used to PCR out GPR-IV and inserted to the linearized CRY2-mCh at its C-terminus using Gibson assembly followed by Dpn1 reaction. CRY2-mCh-AGS3-GPR-IV was PCR amplified using appropriate primers with overhangs containing expected nucleotide mutations to generate CRY2-mCh-GPRcn (1X) consensus. This construct was used as the parent to add subsequent multiple consensus sequences to generate CRY2-mCh-GPRcn(3X) and CRY2-mCh-GPRcn 6X (OptoGDI). To create Lyn-CIBN, we linearized a parent construct with Lyn at the N terminus and incorporated the PCR-amplified CIBN at the C terminus. All cloning was performed using Gibson assembly cloning (NEB). All cDNA constructs were confirmed by sequencing.

### 4.3 Cell culture and transfections

RAW264.7, HeLa, and HEK293T cells were initially purchased from ATCC, USA. Recommended cell culture media: RAW264.7 (RPMI/10%DFBS/1%PS), HeLa (MEM/10%DFBS/1%PS), and HEK293T (DMEM/10%FBS/1%PS) were used to subculture cells at 70-80% (using versene-EDTA (CellGro)) on 29 mm, 60 mm, or 100 mm cell culture dishes in a humidified incubator at 37°C, 5% CO_2_. For live-cell imaging experiments, cells were seeded on 29 mm glass-bottomed dishes at a density of 1 × 10^5^ cells. DNA transfections were performed using either Lipofectamine® 2000 reagent (for HeLa cells) or electroporation (for RAW264.7 cells) according to the manufacturer’s recommended protocols. Briefly, for electroporation of RAW264.7 cells, the following method was used. Nucleofector solution (82 µL), Supplement solution (18 µL) from Amaxa® Cell Line Nucleofector® Kit V, and appropriate volumes of plasmid DNA for specific DNA combinations were mixed. In each electroporation experiment, ∼2 million cells were electroporated using the T-020 method of the Nucleofector™ 2b device (Lonza). Immediately after electroporation, cells were mixed with the cell culture medium at 37 ⁰C. Then, cells were centrifuged at 2800 rpm for 3 minutes. Afterward, the cell pellet was resuspended in appropriate volumes of cell culture media at 37 ⁰C and seeded on glass-bottomed dishes, 200 μL per well. After 2h, 800 μL of more new media was added at 37 ⁰C. Imaging was conducted after ∼5-6 h post-transfection, considering the high expression of constructs. HEK293T cells were plated in a 6-well plate at the density of 2 million cells per well. After 2hrs, cells were treated with 1 mL of complete media and transfected with respective DNA combinations (OptoGDI-1.5 μg, Gα-Rluc8-0.5 μg, Gβ-0.5 μg, and GFP2-Gγ-0.5 μg) with 3.2 μL of PolyJet (SigmaGen) according to manufacturer’s instructions. Then the 6-well plate covered with aluminum foil was kept in a humidified incubator at 37°C, 5% CO_2_ overnight. The next day, transfected cells were washed with PBS once and detached using versine-EDTA (1 mL per well). Detached cells were centrifuged at 200g for 4 min and then resuspended in complete DMEM media. Cells were then seeded at 0.1 million cells per well in 96-well plates coated with poly-D-lysine (100 μg/ mL). Plates were wrapped again with foil and kept in a humidified incubator overnight. Note - (Minimize the light exposure during this process.

### 4.4 Live cell imaging, image analysis, and data processing

The methods, protocols, and parameters for live cell imaging are adapted from previously published work (13, 63, 64). Briefly, live cell imaging experiments were performed using a spinning disk confocal imaging system (Andor Technology) with a 60X, 1.4 NA oil objective, and iXon ULTRA 897BVback-illuminated deep-cooled EMCCD camera. Photoactivation and spatio-temporal light exposure on cells in regions of interest (ROI) was performed using a laser combiner with a 445 nm solid-state laser delivered using Andor® FRAP-PA (fluorescence recovery after photobleaching and photoactivation) unit in real-time, controlled by Andor iQ 3.1 software (Andor Technologies, Belfast, United Kingdom). Fluorescent proteins such as CRY2-mCh, CRY2-mCh-AGS consensus 1X, CRY2-mCh-AGS consensus 3X, OptoGDI, Akt-PH-mCh were imaged using 594 nm excitation−624 nm emission settings. Akt-PH-Venus, Venus-PH, Split Venus β2γ9 were imaged using 515 nm excitation and 542 nm emission. α2AR-CFP was imaged using 445 nm excitation and 478 nm emission, respectively. For global and confined optical activation of CRY2 expressing cells, 445 nm solid-state laser coupled to FRAP-PA was adjusted to deliver 145 nW power at the plane of cells, which scanned light illumination across the region of interest (ROI) at 1 ms/μm^2^. The time-lapse images were analyzed using Andor iQ 3.2 software by acquiring the mean pixel fluorescence intensity changes of the entire cell or selected area/regions of interest (ROIs). Briefly, the background intensity of images was subtracted from the intensities of the ROIs assigned to the desired areas of cells (plasma membrane, endomembranes, and cytosol) before intensity data collection from the time-lapse images. The intensity data from multiple cells were opened in Excel (Microsoft office®) and normalized to the baseline by dividing the whole data set by the average initial stable baseline value. Data were processed further using Origin-pro data analysis software (OriginLab®).

### 4.5 BRET2 assay

After keeping cells in a 96-well plate for overnight, plates were decanted and treated with 90 μL of BRET buffer (20 mM HEPES, 1× HBSS, pH 7.4) per well. Then 10 μL coelenterazine 400a (Nanolight Technology) was added to each well at a final concentration of 2.5 ng/μL After 2 min incubation under dark, plates were read on the Synergy H1 plate reader (Biotek) with 410 nm (RLuc8-coelenterazine 400a) and 515nm (GFP2) emission filters for 1 s integration time per well. After 5 measurements, plates were ejected and exposed to blue light for 30 seconds and continued for another 10 measurements. The GFP to Rluc8 ratio was calculated before and after blue light exposure and the difference in BRET before and after blue light exposure was plotted using Origin-pro data analysis software.

### 4.6 Determination of AGS3 consensus sequence structure and interactions with Gαi via AlphaFold and Schrodinger

We used the amino acid sequence of AGS3 consensus sequence (37) to generate the model protein structure in Fig. 1A using AlphaFold2 (65). The structure that best fits the currently available experimental information was selected for protein folding validation and optimization using the protein preparation tool and loop refinement tool in Schrodinger Bioluminate. We next incorporated a Gαi crystal structure available in PDB database to Schrodinger; we cleaned the structure by removing the unwanted structures, ligands, and solvents and prepared the protein using protein preparation wizard. After that, we docked the prepared proteins using Protein-Protein docking. The best hit with the lowest docking score was selected for further interpretations and to map interactions with Gαi.

### 4.7 Experimental rigor and Statistical analysis

To eliminate potential biases or preconceived notions and improve the experimental rigor, we used the reagent-blinded-experimenter approach for the key findings of our study, i.e., Venus Gγ9 and Split Venus β2γ9 translocation assays. Additionally, OptoGDI-induced cell migration was conducted by two different experimenters. All experiments were repeated multiple times to test the reproducibility of the results. Results are analyzed from multiple cells and represented as mean ± SD. The exact number of cells used in the analysis is given in respective Fig. legends. Digital image analysis was performed using Andor iQ 3.1 software, and fluorescence intensity obtained from regions of interest was normalized to initial values (baseline). Data plot generation and statistical analysis were done using OriginPro software (OriginLab®). One-way ANOVA statistical tests were performed using OriginPro to determine the statistical significance between two or more populations of signaling responses. Tukey’s mean comparison test was performed at the p < 0.05 significance level for the one-way ANOVA statistical test. After obtaining the normalized data, PIP2 recovery rates were calculated using the NonLinear Curve Fitting tool in OriginPro. In the NonLinear Curve Fitting tool, each plot was fitted to DoseResp (dose-response) function under the pharmacology category by selecting the relevant range of data to be fitted. The mean values of hill slopes (P) obtained for each curve fitting are presented as the mean rates of PIP2 recovery.

## Supporting information

Thotamune et al - supporting information

## Author Contribution

W.T. and A.C. made Lyn-CIBN, W.T. performed PIP3 generation, cell migration, and computational modeling experiments and RNAseq and HPA data analysis, S.U. performed PIP2 hydrolysis experiments, C.R. made OptoGDI, performed Gβγ translocation experiments, K.O. made CRY2mCherry-GPRcn(3X), D.K. made CRY2mCherry-GPRcn (1X) and performed Gα specific BRET experiments. D.K., W.T., S.U., C.R., and A.K. conceptualized the project, performed data and statistical analysis, and wrote the original manuscript, W.T., S.U., D.K., C.R., K.O., B.A.C., and A.K. reviewed and finalized the manuscript.

## Data availability

The datasets used and/or analyzed during the current study are available from the corresponding author upon reasonable request.

## Acknowledgment

We acknowledge Dr. N. Gautam (Washington University-School of Medicine, St. Louis, MO, USA) for providing us with plasmid DNA and RNAseq data. We thank the National Eye Institute for providing 11-*cis*-retinal. We thank the Saint Louis University Institute for Drug and Biotherapeutic Innovation for providing computational resources and access to Schrödinger software with funding from the Saint Louis University Research Institute. We thank the Department of Biology at Saint Louis University for various instruments and infrastructure support. We thank Ajith Lab members for all their support in moving the project forward.

