## Supplementary material for "AGS3-based optogenetic GDI induces GPCR-independent Gβγ signaling and macrophage migration": Thotamune et al - supporting information

---

#### Figure S1

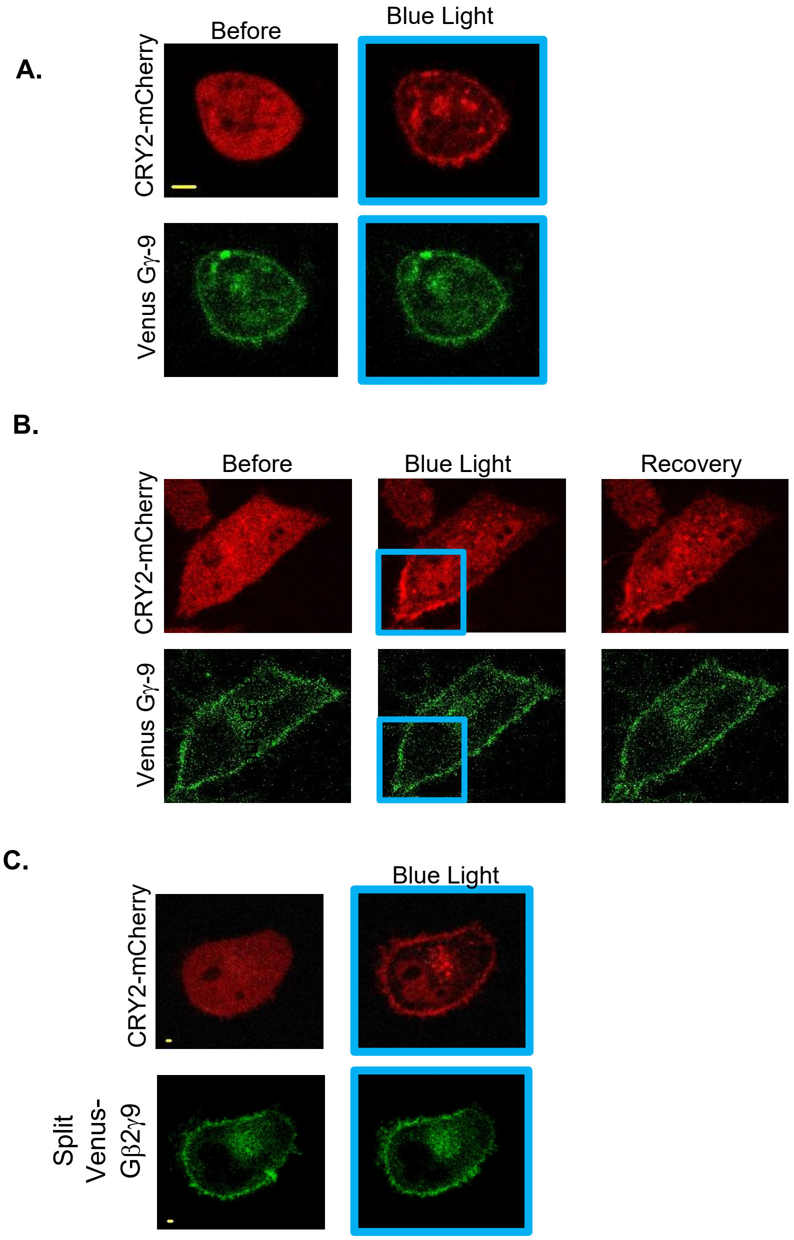

**Figure. S1: (A)** HeLa cells expressing CRY2-mCherry, Lyn-CIBN, and Venus-Gγ9 did not show a detectable Gβγ translocation upon blue light exposure. **(B).** In HeLa cells expressing CRY2-mCherry, Lyn-CIBN and Venus-Gγ9, CRY2-mCherry was recruited to a confined region of the plasma membrane using localized blue light exposure. This subsequent CRY2 recruitment did not induce a detectable subcellular Gβγ translocation. **(C)** In HeLa cells expressing CRY2-mCherry, Lyn-CIBN and split Venus-Gβ2γ9, CRY2-mCherry was recruited to a confined region of the plasma membrane using localized blue light exposure. CRY2-mCherry recruitment did not induce a detectable split Venus-Gβ2γ9 translocation. The scale bar = 5 μm.

### Figure S2

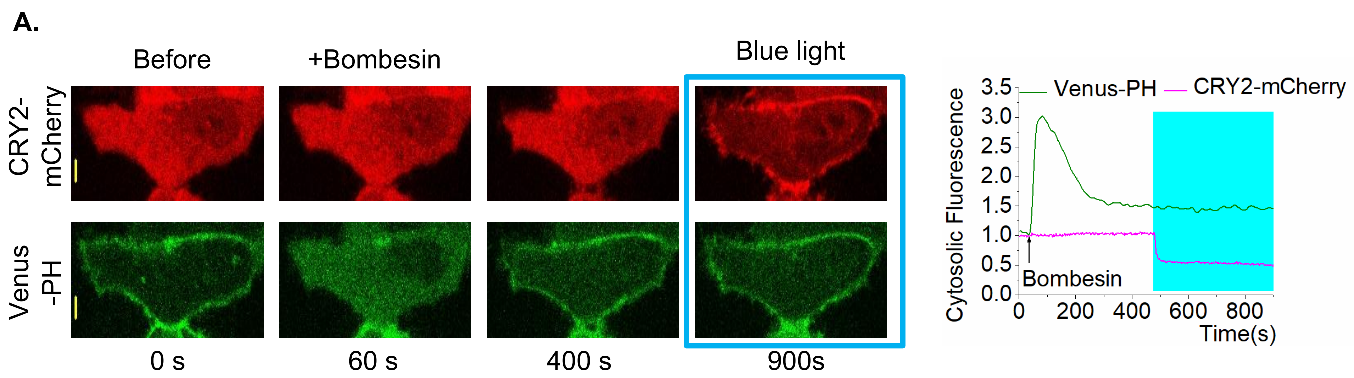


**Figure. S2: (A)** HeLa cells expressing GRPR, Venus-PH, Gβ1, CRY2-mCherry and Lyn-CIBN exhibited robust PIP2 hydrolysis upon 1 μM bombesin addition. CRY2-mCherry was then recruited to the plasma membrane after the PIP2 hydrolysis attenuation reached the equilibrium. The cells did not exhibit a rescue of PIP2 hydrolysis. The scale bar = 5 μm.

#### Figure S3

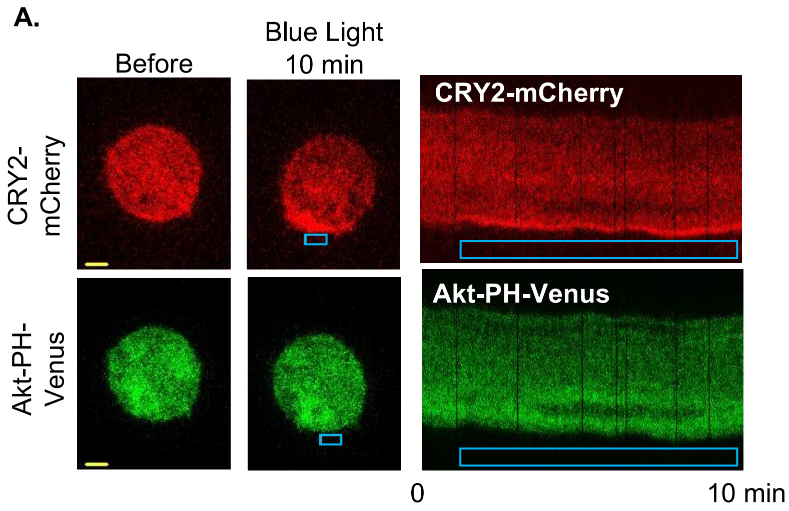


**Figure S3: (A)** RAW cells expressing CRY2-mCherry, Lyn-CIBN, and AKT-PH-Venus were optically activated to recruit Opto-GDI to a confined region (top). The cell does not show localized PIP3 production (bottom images) or detectable migration response towards the blue light. (B) Kymographs of the same cell show the accumulation of Opto-GDI (red) but not Akt-PH-Venus (green) in the leading edge, or the cell migration. The scale bar = 5 µm.

Table S1-A: One-way ANOVA statistics for CRY2-mCh- GPRcn consensus 1X induced Gγ9 translocation extent upon blue light exposure.

| Descriptive Statistics | | | | |
|---|---|---|---|---|
| | N Analysis | Mean | Standard Deviation | SE of Mean |
| Before | 10 | 0.99734 | 0.00943 | 0.00298 |
| After Blue Light | 10 | 1.06374 | 0.02529 | 0.008 |

Table S1-B:

| Overall ANOVA | | | | | |
|---|---|---|---|---|---|
| | DF | Sum of Squares | Mean Square | F Value | Prob>F |
| Model | 1 | 0.02204 | 0.02204 | 60.49522 | <0.0001 |
| Error | 18 | 0.00656 | 3.64321E-4 | | |
| Total | 19 | 0.0286 | | | |

Table S2-A: One-way ANOVA statistics for CRY2-mCh- GPRcn consensus 3X induced Gγ9 translocation extent upon blue light exposure.

| Descriptive Statistics | | | | |
|---|---|---|---|---|
| | N Analysis | Mean | Standard Deviation | SE of Mean |
| Before | 8 | 1.00325 | 0.00965 | 0.00341 |
| After Blue light | 8 | 1.02423 | 0.01654 | 0.00585 |

Table S2-B:

| Overall ANOVA | | | | | |
|---|---|---|---|---|---|
| | DF | Sum of Squares | Mean Square | F Value | Prob>F |
| Model | 1 | 0.00176 | 0.00176 | 9.60401 | 0.00785 |
| Error | 14 | 0.00257 | 1.83395E-4 | | |
| Total | 15 | 0.00433 | | | |

Table S3-A: One-way ANOVA statistics for Opto-GDI induced Gγ9 translocation extent upon blue light exposure.

| Descriptive Statistics | | | | |
|---|---|---|---|---|
| | N Analysis | Mean | Standard Deviation | SE of Mean |
| Before | 9 | 0.99573 | 0.00948 | 0.00316 |
| After Blue Light | 9 | 1.1084 | 0.02912 | 0.00971 |

Table S3-B:

| Overall ANOVA | | | | | |
|---|---|---|---|---|---|
| | DF | Sum of Squares | Mean Square | F Value | Prob>F |
| Model | 1 | 0.05712 | 0.05712 | 121.83117 | <0.0001 |
| Error | 16 | 0.0075 | 4.68862E-4 | | |
| Total | 17 | 0.06462 | | | |

Table S4-A: One-way ANOVA statistics for CRY2-mCh-GPRcn consensus 1X, CRY2-mCh-GPRcn consensus 3X, and Opto-GDI induced Gγ9 translocation extent upon blue light exposure.

| Descriptive Statistics | | | | |
|---|---|---|---|---|
| | N Analysis | Mean | Standard Deviation | SE of Mean |
| CRY2-mCh-GPRcn(1X) | 8 | 1.02423 | 0.01654 | 0.00585 |
| CRY2-mCh-GPRcn(3X) | 10 | 1.06374 | 0.02529 | 0.008 |
| Opto-GDI | 9 | 1.1084 | 0.02912 | 0.00971 |

Table S4-B:

| Overall ANOVA | | | | | |
|---|---|---|---|---|---|
| | DF | Sum of Squares | Mean Square | F Value | Prob>F |
| Model | 2 | 0.03016 | 0.01508 | 25.03591 | <0.0001 |
| Error | 24 | 0.01446 | 6.02373E-4 | | |
| Total | 26 | 0.04462 | | | |

Table S5-A: One-way ANOVA statistics for the PIP2 hydrolysis adaptation rates without Opto-GDI, with cytosolic Opto-GDI and with membrane recruited Opto-GDI

| Descriptive Statistics | | | | |
|---|---|---|---|---|
| | N Analysis | Mean | Standard Deviation | SE of Mean |
| Without Opto-GDI (level 1) | 11 | 0.00812 | 9.79736E-4 | 2.95401E-4 |
| With cytosolic Opto-GDI (level 2) | 9 | 0.00776 | 7.64266E-4 | 2.54755E-4 |
| With membrane recruited opto-GDI (level 3) | 11 | 0.00466 | 4.51568E-4 | 1.36153E-4 |

Table S5-B:

| Overall ANOVA | | | | | |
|---|---|---|---|---|---|
| | DF | Sum of Squares | Mean Square | F value | Prob>F |
| Model | 2 | 7.78551E-5 | 3.89275E-5 | 66.82518 | 2.18846E-11 |
| Error | 28 | 1.63108E-5 | 5.82528E-7 | | |
| Total | 30 | 9.41658E-5 | | | |

Table S6-A: One-way ANOVA statistics for the RAW cell migration distances (μm) with Blue Opsin (GPCR) and Opto-GDI (GPCR independent).

| Descriptive Statistics | | | | |
|---|---|---|---|---|
| | N Analysis | Mean | Standard Deviation | SE of Mean |
| Blue Opsin (GPCR) | 6 | 18.16667 | 4.22453 | 1.72466 |
| Opto-GDI (GPCR independent) | 7 | 7.91429 | 1.65573 | 0.62581 |

Table S6-B:

| Overall ANOVA | | | | | |
|---|---|---|---|---|---|
| | DF | Sum of Squares | Mean Square | F value | Prob>F |
| Model | 1 | 339.5904 | 339.5904 | 35.34659 | <0.0001 |
| Error | 11 | 105.6819 | 9.60745 | | |
| Total | 12 | 445.27231 | | | |

Table S7-A: One-way ANOVA statistics for the RAW cell migration distances (μm) with Opto-GDI (GPCR independent) and Opto-GDI (GPCR independent) with Wortmannin (50 nM).

| Descriptive Statistics | | | | |
|---|---|---|---|---|
| | N Analysis | Mean | Standard Deviation | SE of Mean |
| Opto-GDI (GPCR independent) | 7 | 7.91429 | 1.65573 | 0.62581 |
| Opto-GDI (GPCR independent) + Wortmannin (50 nM) | 6 | 0.36667 | 0.27325 | 0.11155 |

Table S7-B:

| Overall ANOVA | | | | | |
|---|---|---|---|---|---|
| | DF | Sum of Squares | Mean Square | F value | Prob>F |
| Model | 1 | 184.04579 | 184.04579 | 120.34925 | <0.0001 |
| Error | 11 | 16.8219 | 1.52926 | | |
| Total | 12 | 200.86769 | | | |

Table S8-A: One-way ANOVA statistics for the RAW cell migration distances (μm) with Opto-GDI (GPCR independent) and Opto-GDI (GPCR independent) with Gallein (10 μM).

| Descriptive Statistics | | | | |
|---|---|---|---|---|
| | N Analysis | Mean | Standard Deviation | SE of Mean |
| Opto-GDI (GPCR independent) | 7 | 7.91429 | 1.65573 | 0.62581 |
| Opto-GDI (GPCR independent) + Gallein (10 μM) | 7 | 2.15714 | 0.70677 | 0.26713 |

Table S8-B:

| Overall ANOVA | | | | | |
|---|---|---|---|---|---|
| | DF | Sum of Squares | Mean Square | F value | Prob>F |
| Model | 1 | 116.00643 | 116.00643 | 71.58786 | <0.0001 |
| Error | 12 | 19.44571 | 1.62048 | | |
| Total | 13 | 135.45214 | | | |

Table S9-A: One-way ANOVA statistics for the RAW cell migration velocities (μm/s) with Blue Opsin (GPCR) and Opto-GDI (GPCR independent).

| Descriptive Statistics | | | | |
|---|---|---|---|---|
| | N Analysis | Mean | Standard Deviation | SE of Mean |
| Blue Opsin (GPCR) | 6 | 0.03028 | 0.00704 | 0.00287 |
| Opto-GDI (GPCR independent) | 7 | 0.01319 | 0.00276 | 0.00104 |

Table S9-B:

| Overall ANOVA | | | | | |
|---|---|---|---|---|---|
| | DF | Sum of Squares | Mean Square | F value | Prob>F |
| Model | 1 | 9.43307E-4 | 9.43307E-4 | 35.34659 | <0.00011 |
| Error | 11 | 2.93561E-4 | 2.66873E-5 | | |
| Total | 12 | 0.00124 | | | |

Table S10-A: One-way ANOVA statistics for the RAW cell migration velocities (μm/s) with Blue Opsin (GPCR) and Opto-GDI (GPCR independent) with Wortmannin (50 nM).

| Descriptive Statistics | | | | |
|---|---|---|---|---|
| | N Analysis | Mean | Standard Deviation | SE of Mean |
| Opto-GDI (GPCR independent) | 7 | 0.01319 | 0.00276 | 0.00104 |
| Opto-GDI (GPCR independent) + Wortmannin (50 nM) | 6 | 6.11111E-4 | 4.5542E-4 | 1.85924E-4 |

Table S10-B:

| Overall ANOVA | | | | | |
|---|---|---|---|---|---|
| | DF | Sum of Squares | Mean Square | F value | Prob>F |
| Model | 1 | 5.11238E-4 | 5.11238E-4 | 120.34925 | <0.0001 |
| Error | 11 | 4.67275E-5 | 4.24796E-6 | | |
| Total | 12 | 5.57966E-4 | | | |

Table S11-A: One-way ANOVA statistics for the RAW cell migration velocities (μm/s) with Blue Opsin (GPCR) and Opto-GDI (GPCR independent) with Gallein (10 μM).

| Descriptive Statistics | | | | |
|---|---|---|---|---|
| | N Analysis | Mean | Standard Deviation | SE of Mean |
| Opto-GDI (GPCR independent) | 7 | 0.01319 | 0.00276 | 0.00104 |
| Opto-GDI (GPCR independent) + Gallein (10 μM) | 7 | 0.0036 | 0.00118 | 4.45223E-4 |

Table S11-B:

| Overall ANOVA | | | | | |
|---|---|---|---|---|---|
| | DF | Sum of Squares | Mean Square | F value | Prob>F |
| Model | 1 | 3.2224E-4 | 3.2224E-4 | 71.58786 | <0.0001 |
| Error | 12 | 5.40159E-5 | 4.50132E-6 | | |
| Total | 13 | 3.76256E-4 | | | |

**Movie S1.** Blue light induced OptoGDI recruitment in a HeLa cell expressing OptoGDI, Lyn-CIBN and Venus-Gγ9.

**Movie S2.** HeLa cell expressing OptoGDI, Lyn-CIBN and Venus-Gγ9 shows Venus-Gγ9 translocation upon blue light induced OptoGDI recruitment.

**Movie S3.** Blue light induced Opto-GDI recruitment to the plasma membrane in a HeLa cell expressing GRPR, OptoGDI, Lyn-CIBN and Venus-PH.

**Movie S4.** Blue light + Opto-GDI recruitment induced PIP2 hydrolysis rescue in a HeLa cell expressing GRPR, OptoGDI, Lyn-CIBN and Venus-PH. The movie shows the PIP2 sensor, Venus-PH dynamics in the cytosol upon bombesin addition and OptoGDI recruitment.

**Movie S5.** A Raw264.7 cell expressing Opto-GDI (mCherry-tagged), Lyn-CIBN, and Akt-PH-Venus shows Opto-GDI recruitment to the blue light exposed side of the cells. The box shows blue light exposure (1 Hz) during imaging. Note the migration of the cell towards blue light.

**Movie S6.** The same Raw264.7 cell expressing Opto-GDI (mCherry-tagged), Lyn-CIBN, and Akt-PH-Venus shows PIP3 generation at the Opto-GDI recruited, blue light exposed side of the cells. The box shows blue light exposure (1 Hz) during imaging. Note the migration of the cell towards blue light.

**Movie S7.** Raw264.7 cell expressing blue opsin and Akt-PH-mCherry shows PIP3 generation at the blue light exposed side of the cell. The box shows blue light exposure (1 Hz) during imaging. Note the migration of the cell towards blue light.
